# Nuclear Factor I in neurons, glia and during the formation of Müller glia-derived progenitor cells in avian, porcine and primate retinas

**DOI:** 10.1101/2021.07.13.451621

**Authors:** Heithem M. El-Hodiri, Warren A. Campbell, Lisa E. Kelly, Evan C. Hawthorn, Maura Schwartz, Archana Jalligampala, Maureen A. McCall, Kathrin Meyer, Andy J. Fischer

## Abstract

The regenerative potential of Müller glia (MG) is extraordinary in fish, poor in chick and terrible in mammals. In the chick model, MG readily reprogram into proliferating Müller glia-derived progenitor cells (MGPCs), but neuronal differentiation is very limited. The factors that suppress the neurogenic potential of MGPCs in the chick are slowly being revealed. Isoforms of Nuclear Factor I (NFI) are cell-intrinsic factors that limit neurogenic potential; these factors are required for the formation of MG in the developing mouse retina (Clark et al., 2019) and deletion of these factors reprograms MG into neuron-like cells in mature mouse retina (Hoang et al., 2020). Accordingly, we sought to characterize the patterns of expression NFIs in the developing, mature and damaged chick retina. In addition, we characterized patterns of expression of NFIs in the retinas of large mammals, pigs and monkeys. Using a combination of single cell RNA-sequencing (scRNA-seq) and immunolabeling we probed for patterns of expression. In embryonic chick, levels of NFIs are very low in early E5 (embryonic day 5) retinal progenitor cells (RPCs), up-regulated in E8 RPCs, further up-regulated in differentiating MG at E12 and E15. NFIs are maintained in mature resting MG, microglia and neurons. Levels of NFIs are reduced in activated MG in retinas treated with NMDA and/or insulin+FGF2, and further down-regulated in proliferating MGPCs. However, levels of NFIs in MGPCs were significantly higher than those seen in RPCs. Immunolabeling for NFIA and NFIB closely matched patterns of expression revealed in different types of retinal neurons and glia, consistent with findings from scRNA-seq. In addition, we find expression of NFIA and NFIB through progenitors in the circumferential marginal zone at the far periphery of the retina. We find similar patterns of expression for NFIs in scRNA-seq databases for pig and monkey retinas. Patterns of expression of NFIA and NFIB were validated with immunofluorescence in pig and monkey retinas wherein these factors were predominantly detected in MG and a few types of inner retinal neurons. In summary, NFIA and NFIB are prominently expressed in developing chick retina and by mature neurons and glia in the retinas of chicks, pigs and monkeys. Although levels of NFIs are decreased in chick, in MGPCs these levels remain higher than those seen in neurogenic RPCs. We propose that the neurogenic potential of MGPCs in the chick retina is suppressed by NFIs.

## Introduction

Müller glia (MG) are the primary glial cell of the retina that serves pleiotropic functions including neurotransmitter recycling, osmotic balance, structural support, potassium siphoning, and bicarbonate transport (Reichenbach and Bringmann, 2013). Importantly, MG retain stem cell properties and are capable of replacing lost neurons. However, the regenerative potential of MG is variable among species. Zebrafish can regenerate all retinal neurons, while mice fail to form any MG-derived progenitor cells (MGPCs). MG in chick retinas have an intermediate potential with significant proliferation of MGPCs but limited neurogenesis into a few neuronal subtypes. The limited regenerative potential in chicks serves as an insightful *in vivo* model to investigate factors that influence retinal regeneration. The factors underlying the limited neurogenic capacity of MGPCs in the chick retina remain poorly understood.

Nuclear Factor I (NFI) gene products are known to promote gliogenesis (Deneen et al., 2006; Wilczynska et al., 2009). Studies in the developing nervous system have indicated that the loss-of-function mutations of NFI genes suppress glial differentiation (Namihira et al., 2009; Piper et al., 2010; Kang et al., 2012). NFIs, including NFIA, NFIB, NFIC and NFIX, are known to play important roles in the development of glia in the central nervous system. Further, NFIs are required for the differentiation of MG in the developing rodent retina (Clark et al., 2019) and the targeted knock-out of *Nfia*, *Nfib* and *Nfix* promotes the reprogramming of MG into neurons in adult damaged mouse retinas (Hoang et al., 2020). A comparative genomic analysis has revealed that NFIs are key components of regulatory networks that maintain resting glial phenotype of mouse MG and these regulatory “hubs” are absent in zebrafish MG (Hoang et al., 2020).

In zebrafish, the pathways that drive the initial de-differentiation and reprogramming of MG include MAPK-ERK, Gsk3β-β catenin, and Jak-stat. Similarly, factors such as let-7, TGFβ, and insulinoma-associated 1a (Insm1a) drive progenitors out of the cell cycle to enable neurogenesis (Goldman, 2014). Many of these important cell signaling pathways can be recapitulated in the chick retina to stimulate MGPC formation and enhance neurogenesis (Todd and Fischer, 2015; Todd et al., 2016a; Zelinka et al., 2016; Todd et al., 2017a, 2018). However, neurogenesis from MGPCs in the chick retina remains very limited suggesting that there may be cell intrinsic factors that suppress neurogenic potential. Cell signaling has been shown to influence from neurogenesis from MGPCs. In the chick, neuronal differentiation of the progeny of MGPCs can be enhanced by inhibiting Notch (Hayes et al., 2007; Ghai et al., 2010), inhibiting glucocortioids (Gallina et al., 2014), activation of retinoic acid (Todd et al., 2018), and inhibition of Jak/Stat-signaling (Todd et al., 2016a). In the mouse retina, neuronal differentiation from Ascl1a-overexpressing MG can be enhanced by inhibiting Jak/Stat-signaling (Jorstad et al., 2020) or ablating reactive microglia (Todd et al., 2020).

During neural development there are many factors that act to suppress neurogenesis and in favor of gliogenesis. For example, Notch-signaling and downstream transcription effector Hes1/Hes5 have been shown to promote glial specification and differentiation during later stages of retinal development (Furukawa et al., 2000; Vetter and Moore, 2001; Bernardos et al., 2005). During late stages of development, TGFB suppresses the proliferation of progenitors and maturing MG in the rat retina (Close et al., 2005). In addition, TGFB/Smad2/3-signaling suppresses the formation of MGPCs in chick and zebrafish retinas (Lenkowski et al., 2013; Tappeiner et al., 2016; Todd et al., 2017a; Lee et al., 2020) and act to coordinate activity between microglia and MG (Ma et al., 2019). During mouse retinal development knock-out *Nfia, Nfib* and *Nfix* results in a loss of MG and a prolonged period of proliferation postnatally (Clark et al., 2019). In the adult mouse retina, knock-out *Nfia, Nfib* and *Nfix* specifically in MG when combined with NMDA-induced damage and insulin+FGF2 results in the generation of cells that resemble inner retinal neurons (Hoang et al., 2020).

The purpose of this study was to characterize patterns of expression of NFIs in the chick retina during developing, in the mature retina, and following treatments that induce the formation of proliferating MGPCs. Further, we analyze patterns of expression of NFIs in the retinas of large mammals, pig and non-human primate.

## Methods and Materials

### Animals

#### Avian

Fertilized eggs were obtained from the Michigan State University, Department of Animal Science. Eggs were incubated at 37.5°C, with a 1hr cool-down to room temperature every 24hrs, rotated every 45 minutes, and maintained at a humidity of 45%. Embryos were harvested at different time points after incubation and staged according to guidelines established by Hamburger and Hamilton (Hamburger, 1951). The use of hatched chicks followed guidelines established by the National Institutes of Health and IACUC at the Ohio State University. P0 wildtype leghorn chicks (*Gallus gallus domesticus*) were obtained from Meyer Hatchery (Polk, Ohio). Post-hatch chicks were housed in stainless-steel brooders at 25°C with a diurnal cycle of 12 hours light, 12 hours dark (8:00 AM-8:00 PM) and provided water and Purina^tm^ chick starter *ad libitum*.

#### Porcine

Wild-type pigs born from a heterozygous cross of a transgenic P23H miniature pig model were utilized for this study. The pigs were housed at the Kentucky Lions Eye Institute (Louisville, Kentucky) and managed according to the guidelines of the ARVO Statement for the Use of Animals in Ophthalmic and Vision Research. Pigs were fed a swine-specific feed (Harlan Teklad Miniswine Diet 8753; Envigo, Hackensack, NJ) twice daily and provided water *ad libitum*. Pigs were kept on a 14/10- hour light/dark cycle.

#### Primate

Cynomologous macaques (*Macaca fascicularis*) were housed at the Mannheimer Foundation (Homestead, Florida). Regular monitoring of overall health and body weight was performed to assess the welfare of the animals. All procedures were performed in accordance with the National Institutes of Health and approved by IACUC at the Mannheimer Foundation.

### Intraocular injections

Chicks were anesthetized with 2.5% isoflurane mixed with oxygen from a non-rebreathing vaporizer. The intraocular injections were performed as previously described (Fischer et al., 1998). With all injection paradigms, both pharmacological and vehicle treatments were administered to the right and left eye respectively. Compounds were injected in 20 μl sterile saline with 0.05 mg/ml bovine serum albumin added as a carrier. Compounds included: NMDA (500 nmol/dose; Sigma-Aldrich).

### Preparation of clodronate liposomes

Clodronate liposomes were synthesized utilizing a modified protocol from previous descriptions (Van Rooijen, 1989; van Rooijen, 1992; Zelinka et al., 2012). In short, approximately 8 mg of L-α-Phosphatidyl-DL-glycerol sodium salt (Sigma P8318) was dissolved in chloroform. 50 mg of cholesterol was dissolved in chloroform with the lipids in a sterile microcentrifuge tube. This tube was rotated under nitrogen gas to evaporate the chloroform and leave a thin lipid-film on the walls of the tube. 158 mg dichloro-methylene diphosphonate (clodronate; Sigma-Aldrich) dissolved in sterile PBS (pH 7.4) was added to the lipid/cholesterol film and vortexed for 5 minutes. To reduce size variability of lipid vesicles, the mixture was sonicated at 42 kHz for 6 minutes.

Purification of liposomes was accomplished via centrifugation at 10,000 x G for 15 minutes, aspirated, and resuspended in 150 µl PBS. Each retinal injection used between 5 and 20 ul of clodronate-liposome solution. There was a variable yield of clodronate-liposomes during the purification resulting in some variability per dose. The dosage was adjusted such that >98% of the microglia are ablated by 2 days after administration with no off-target cell death or pigmented epithelial cells.

### Single Cell RNA sequencing of retinas

Retinas were obtained from embryonic chicks, hatched chicks, adult pigs and adult macaque monkeys. Isolated chick and macaque retinas were dissociated in a 0.25% papain solution in Hank’s balanced salt solution (HBSS), pH = 7.4, for 20-40 minutes, and suspensions were frequently triturated. Pig retinas were dissociated similarly except that papain was diluted using Earle’s balanced salt solution (EBSS). The dissociated cells were passed through a sterile 70µm filter to remove large particulate debris. Dissociated cells were assessed for viability (Countess II; Invitrogen) and cell-density diluted to 700 cell/µl. Each single cell cDNA library was prepared for a target of 10,000 cells per sample. The cell suspension and Chromium Single Cell 3’ V2 or V3 reagents (10X Genomics) were loaded onto chips to capture individual cells with individual gel beads in emulsion (GEMs) using 10X Chromium Controller. cDNA and library amplification for an optimal signal was 12 and 10 cycles respectively.

Sequencing was conducted on Illumina HiSeq4000 (Novogene) with 30 bp for Read 1 and 98 bp for Read 2. Fasta sequence files were de-multiplexed, aligned, and annotated using the appropriate ENSMBL database (GRCg6a for chicken, Sscrofa 11.1 for pig, Mmul_10 for macaque) and Cell Ranger software. Gene expression was counted using unique molecular identifier bar codes, and gene-cell matrices were constructed. Using Seurat toolkits, Uniform Manifold Approximation and Projection for Dimension Reduction (UMAP) plots were generated from aggregates of multiple scRNA-seq libraries (Satija et al., 2015; Butler et al., 2018). Seurat V3 was used to construct violin/scatter plots. Significance of difference in violin/scatter plots was determined using a Wilcoxon Rank Sum test with Bonferroni correction. Monocle was used to construct unbiased pseudo-time trajectories and scatter plotters for MG and MGPCs across pseudotime (Trapnell et al., 2012; Qiu et al., 2017a; b). Genes that were used to identify different types of retinal cells included the following: (1) Müller glia: *GLUL, VIM, SCL1A3, RLBP1*, (2) MGPCs: *PCNA, CDK1, TOP2A, ASCL1*, (3) microglia: *C1QA, C1QB, CCL4, CSF1R, TMEM22*, (4) ganglion cells: *THY1, POU4F2, RBPMS2, NEFL, NEFM*, (5) amacrine cells: *GAD67, CALB2, TFAP2A*, (6) horizontal cells:

*PROX1, CALB2, NTRK1*, (7) bipolar cells: *VSX1, OTX2, GRIK1, GABRA1*, and (7) cone photoreceptors: *CALB1, GNAT2, OPN1LW*, and (8) rod photoreceptors: *RHO, NR2E3, ARR3.* The MG have an over-abundant representation in the scRNA-seq databases.

This likely resulted from fortuitous capture-bias and/or tolerance of the MG to the dissociation process.

### Fixation, sectioning and immunocytochemistry

Retinal tissue samples were formaldehyde fixed, sectioned, and labeled via immunohistochemistry as described previously (Fischer et al., 2008; Ghai et al., 2009). Samples processed for immunohistochemistry using antibodies against NFIB were fixed for a shorter period of time (10 minutes) and were pre-treated with 1% (w/v) saponin (Sigma-Aldrich) in phosphate-buffered saline (PBS; Fisher Scientific). Antibody dilutions and commercial sources for images used in this study are described in table 1.

**Table 1.** List of antibodies, working dilution, clone/catalog number and source.

| Antibody | Dilution | Host | Clone/Catalog number | Source |
| --- | --- | --- | --- | --- |
| NFIA | 1:300 | Rabbit | HPA006111 | Atlas Antibodies |
| NFIB | 1:300 | Rabbit | LS-B13531 | LSBio |
| Sox2 | 1:1000 | Goat | KOY0418121 | R&D |
| Sox9 | 1:2000 | Rabbit | AB5535 | Millipore |
| CD45 | 1:200 | Mouse | HIS-C7 | Prionics |
| Nkx2.2 | 1:100 | Mouse | 74.5A5 | DSHB |
| Neurofilament | 1:100 | Mouse | RT97 | DSHB |
| AP2-alpha | 1:50 | Mouse | AP2A | DSHB |
| Brn3a | 1:50 | Mouse | MAB1585 | Chemicon |
| Islet1 | 1:50 | Mouse | 40.2D6 | DSHB |
| Otx2 | 1:1000 | Goat | AF1979 | RD Systems |
| PCNA | 1:1000 | Mouse | M0879 | Dako |
| p21-CIP1 (CDKN1A) | 1:200 | Rabbit | B9242 | LSBio |

The antisera used in this study included: (**1**) rabbit anti-NFIA was used at 1:300 (HPA006111; Atlas Antibodies). This antibody was raised to Recombinant Protein Epitope Signature Tag (PrEST) amino acid LKSVEDEMDSPGEEPFYTGQGRSPGS GSQSSGWHEVEPGMPSPTTLKKSEKSGFSSPSPSQTSSLGTAFTQHHRPVITGPRASPHATPSTLHFPTSPIIQQ. Patterns of immunolabeling for NFIA precisely match patterns of expression of *NFIA* in scRNA-seq databases for retinal cells from chicks, pigs and monkeys (current study). (Antibody Registry ID: AB_185442). (**2**) rabbit anti-NFIB was used at 1:300 (LS-B13531; LSBio). This antibody was raised to amino acids 325 and 375 of human Nuclear Factor I/B. This antibody was validated by IHC on formalin fixed paraffin embedded human heart and lung and by western blot (manufacturer). Patterns of immunolabeling for NFIB precisely match patterns of expression of *NFIB* in scRNA-seq databases for retinal cells from chicks, pigs and monkeys (current study). (Antibody Registry ID: AB_185442). (**3**) goat anti-Sox2 was used at 1:1000 (Y-17; Santa Cruz Biotechnology). The antibody was raised to the recombinant C-terminus of human Sox2 and recognizes a single 34 kDa band in Western blot analysis of lysate from mouse embryonic stem cells (manufacturer). The Sox2 antibody is known to recognize amino acids 277-293 of human Sox2, as determined by preabsorption controls and mass spectrometry analysis of blocking peptide (Poche et al., 2008). The Sox2 antibody produced a pattern of labeling in the nuclei of progenitors, Müller glia, astrocytes and cholinergic amacrine cells in the retina of different vertebrate species, consistent with previous reports (Fischer et al., 2010b). Patterns of immunolabeling for Sox2 precisely match patterns of expression of *SOX2* in scRNA-seq databases for retinal cells from chicks, pigs and monkeys (Supplemental Fig. 1). (Antibody Registry ID: AB_2286684). (4) rabbit anti-Sox9 was used at 1:2000 (AB5535; Millipore). This antibody was raised against human synthetic peptide amino acids VPSIPQTHSPQWEQPVYTQLTRP. Rabbit anti-Sox9 detects a 60-65 kDa band on Western blot analysis of mouse brain tissue (manufacturer’s information). *In situ* hybridization analysis of Sox9 mRNA in the embryonic retina produces an identical pattern to that seen using the Sox9 antibody (Wright et al., 1995; Poche et al., 2008). (Antibody Registry ID: AB_2239761). (5) mouse anti-CD45 was used at 1:200 (HIS-C7; Prionics). This antibody was raised to chicken CD45 HIS-C7 monoclonal antibody (diluted 1/40; Cedi Diagnostics, Lelystad, The Netherlands; catalog 7500970, lot 03M020), obtained from the hybridoma cell line Ch/CD 45 HIS-C7, which is a specific marker for macrophages/microglial cells in the developing and mature chick central nervous system (Cuadros et al., 2006; Brunet et al., 2007). (Antibody Registry ID: AB_2314144). (6) mouse anti-Nkx2.2 was used at 1:50 (74.5A5, Developmental Studies Hybridoma Bank; DSHB). This antibody was raised to Nkx2.2-GST fusion protein expressed in *E. coli*. Patterns of immunolabeling for Nkx2.2 have been well-established in in chick retina (Fischer et al., 2010b; a) and precisely match patterns of expression of *NKX2-2* in scRNA-seq databases of chick retina (Campbell et al., 2019, 2021). (Antibody Registry ID: AB_531794). (**7**) mouse anti-neurofilament was used at 1:50 (RT97, DSHB). This antibody was raised to semi-purified neurofilaments from rat brain homogenate and recognizes phospho-epitopes on the tail domain of neurofilament at KSPXK or KSPXXXK motifs. Patterns of immunolabeling for neurofilament precisely match expression patterns observed with ISH (Fischer et al., 2004) and scRNA-seq from chick retina (Supplemental Fig. 1). (Antibody Registry ID: AB_528399). (**8**) mouse anti-AP2-alpha was used at 1:50 (AP2A; DSHB). This antibody was raised to full length recombinant human AP2-alpha. Patterns of immunolabeling for Nkx2.2 have been well- established in in chick retina (Fischer et al., 2007; Campbell et al., 2021) and precisely match patterns of expression of *TFAP2A* in scRNA-seq databases of chick, pig and monkey retinas (Supplemental Fig. 1). (Antibody Registry ID: AB_2619181). (**9**) mouse anti-Brn3a was used at 1:50 (MAB1585, clone 5A3.2 Millipore). This antibody was raised to amino acids 186-224 of Brn-3a fused to the T7 gene 10 protein (Xiang et al., 1995). This antibody shows no reactivity to Brn-3a knock-out mice (Xiang et al., 1996). Patterns of immunolabeling for Brn3a-alpha precisely match patterns of expression of *POU4F2* in scRNA-seq databases of chick retina (Gallina et al., 2014). (Antibody Registry ID: AB_94166). (**10**) mouse anti-Islet1 was used at 1:50 (40.2D6; DSHB). This antibody was raised to the C-terminus (amino acids 247–349) of rat Islet1. The 40.2D6 monoclonal antibody is known to recognize both Islet1 and Islet2 and has a well-established pattern of labeling in the chicken retina (Fischer et al., 2008). Patterns of immunolabeling for Islet1 precisely match patterns of expression of *ISL1* in scRNA-seq databases of chick retina (Supplemental Fig. 1). (Antibody Registry ID: AB_528315). (**11**) goat anti-Otx2 was used at 1:1000 (AF1979; R&D Systems). This antibody was raised to recombinant human Otx2 (Met1-Leu289). Validation of this antibody has been performed via Western blot shows lysates of IMR-32 human neuroblastoma cell line (manufacturer). Patterns of immunolabeling for Otx2 precisely match patterns of expression of *OTX2* in scRNA-seq databases of chick, pig and monkey (Supplemental Fig. 1). (Antibody Registry ID: AB_2157172). (**12**) mouse anti-PCNA was used at 1:1000 (M0879; Dako Immunochemicals). This antibody was raised to a synthetic amino acid sequence (LVFEAPNQEK) from rat PCNA. Mouse anti-PCNA recognizes a single band at ∼36 kDa on western blot analysis of rat retinal extract (Gordon et al., 2002). This antibody has been shown to recognize proliferating cells in the rodent retina (Sigulinsky et al., 2008) chick retina (Fischer and Reh, 2000; Ghai et al., 2008). Patterns of immunolabeling for PCNA precisely match patterns of expression of *PCNA* in scRNA-seq databases of chick retina (Supplemental Fig. 1d). (Antibody Registry ID: AB_2160651). (**13**) rabbit anti-p21-CIP1 was used at 1:200 (B9242 LSBio) This antibody was raised to a synthetic peptide (amino acids 111-160) from human p21- Cip1. Validation for IHC was done on formalin fixed paraffin embedded human uterus and endometrium tissue and peptide ELISA. Western blot with rabbit anti-p21-CIP1 recognizes a ∼26 kDa on western blot of COLO and K562 cells, and absent when blocked with the synthesized peptide. Patterns of immunolabeling for p21-CIP1 precisely match patterns of expression of *CDKN1A* in scRNA-seq databases of chick retina (Supplemental Fig. 1).

**Figure 1.**
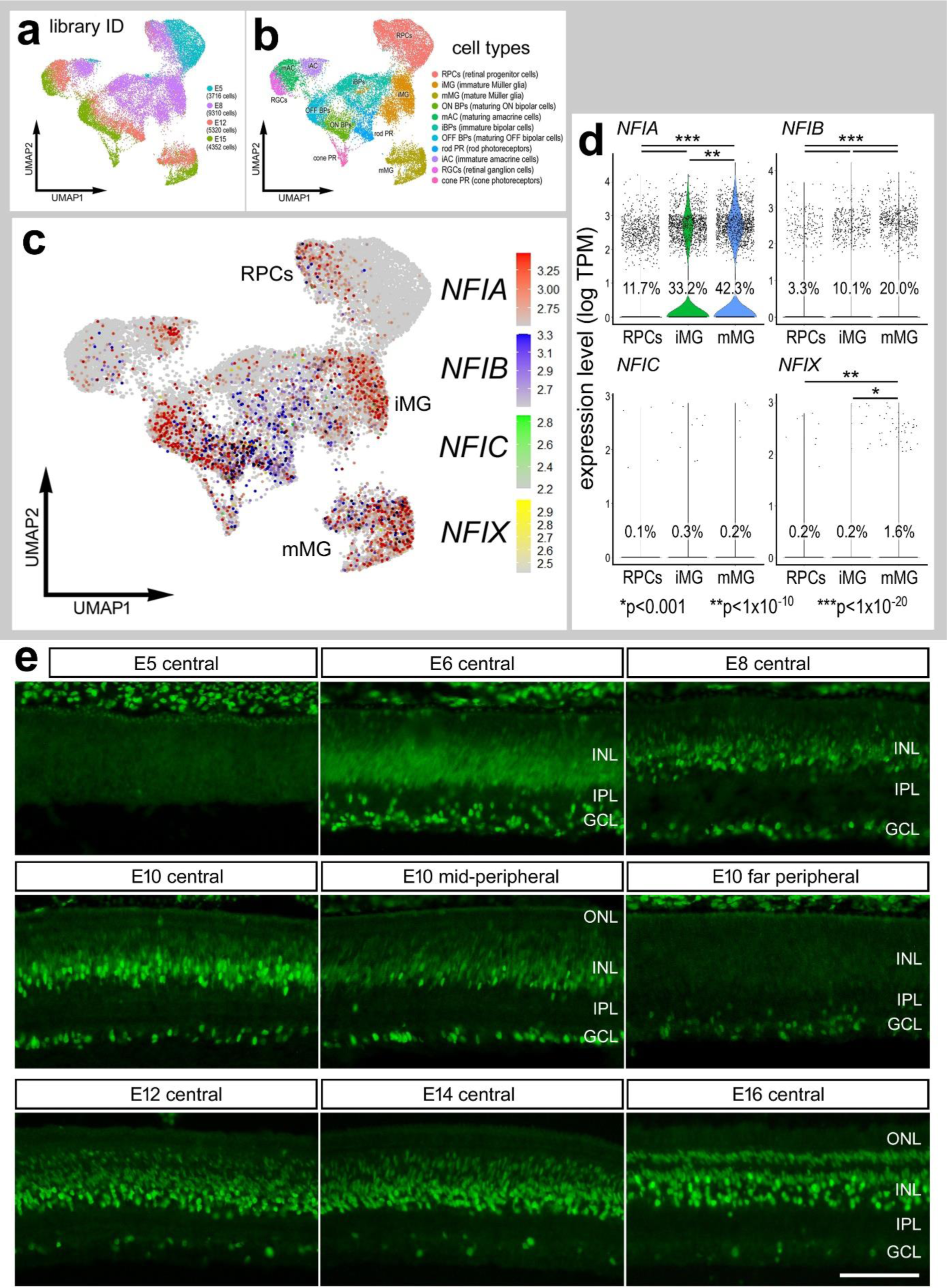
Expression of NFIs in developing chick retinas. scRNA-seq was used to identify patterns of expression of NFIs among embryonic retinal cells at four stages of development, E5, E8, E12 and E15. UMAP-ordered clusters of cells were identified based on expression of cell-distinguishing markers (**a,b**). A heatmap of *NFIA, NFIB, NFIC* and *NFIX* illustrates expression profiles in different developing retinal cells (**c**). Each dot represents one cell and black dots indicate cells with 2 or more genes expressed. Violin plots illustrate the upregulation of *NFIA*, *NFIB* and *NFIX* in maturing MG (**d**). The number on each violin indicates the percentage of expressing cells. Significant difference (*p<0.01, **p<0.0001, ***p<<0.0001) in levels of expression was determined by using a Wilcox rank sum with Bonferoni correction. (**e**) Antibodies to NFIA were applied to vertical sections of central and peripheral regions of retinas from embryos at E5, E6, E8, E10, E12, E14 and E16. The calibration bar in **e** represents 50 µm. RPC – retinal progenitor cell, MG – Müller glia, iMG – immature Müller glia, mMG - mature Müller glia, ONL – outer nuclear layer, INL – inner nuclear layer, IPL – inner plexiform layer, GCL – ganglion cell layer.

Observed labeling was not due to off-target labeling of secondary antibodies or tissue autofluorescence because sections incubated exclusively with secondary antibodies were devoid of fluorescence. Secondary antibodies utilized include donkey-anti-goat-Alexa488/568, goat-anti-rabbit-Alexa488/568/647, goat-anti-mouse- Alexa488/568/647, goat-anti-rat-Alexa488 (Life Technologies) diluted to 1:1000 in PBS and 0.1% Triton X-100.

### Photography, measurements, cell counts and statistics

Microscopy images of retinal sections were captured with the Leica DM5000B microscope with epifluorescence and the Leica DC500 digital camera. High resolution confocal images were obtained with a Leica SP8 available in the Department of Neuroscience Imaging Facility at The Ohio State University. Representative images were modified to enhance brightness and contrast using Adobe Photoshop. The retinal region selected for investigation was standardized between treatment and control groups to reduce variability and improve reproducibility.

## Results

### *NFIs* in embryonic retina

We re-probed scRNA-seq databases that have previously been used to compare MG and MGPCs across fish, chick and mouse model systems (Hoang et al., 2020) and characterize expression patterns of genes related to NFkB-signaling (Palazzo et al., 2020), midkine-signaling (Campbell et al., 2021) and matrix metalloproteases (Campbell et al., 2019). scRNA-seq libraries were established for retinas from chick embryos at E5, E8, E12, and E15. The aggregate library contained 22,698 cells after excluding doublets, and cells with low UMI and <400 genes. UMAP plots of embryonic retinal cells formed clustered of cells that correlated with developmental stage and cell type (Fig. 1a,b). Cell types were identified based on expression of well-established markers. Specifically, retinal progenitor cells from E5 and E8 retinas were identified by expression of *ASCL1*, C*DK1*, and *TOP2A*, and maturing MG were identified by expression of *GLUL*, *RLBP1* and *SLC1A3* (Supplemental Fig. 2a,b).

**Figure 2.**
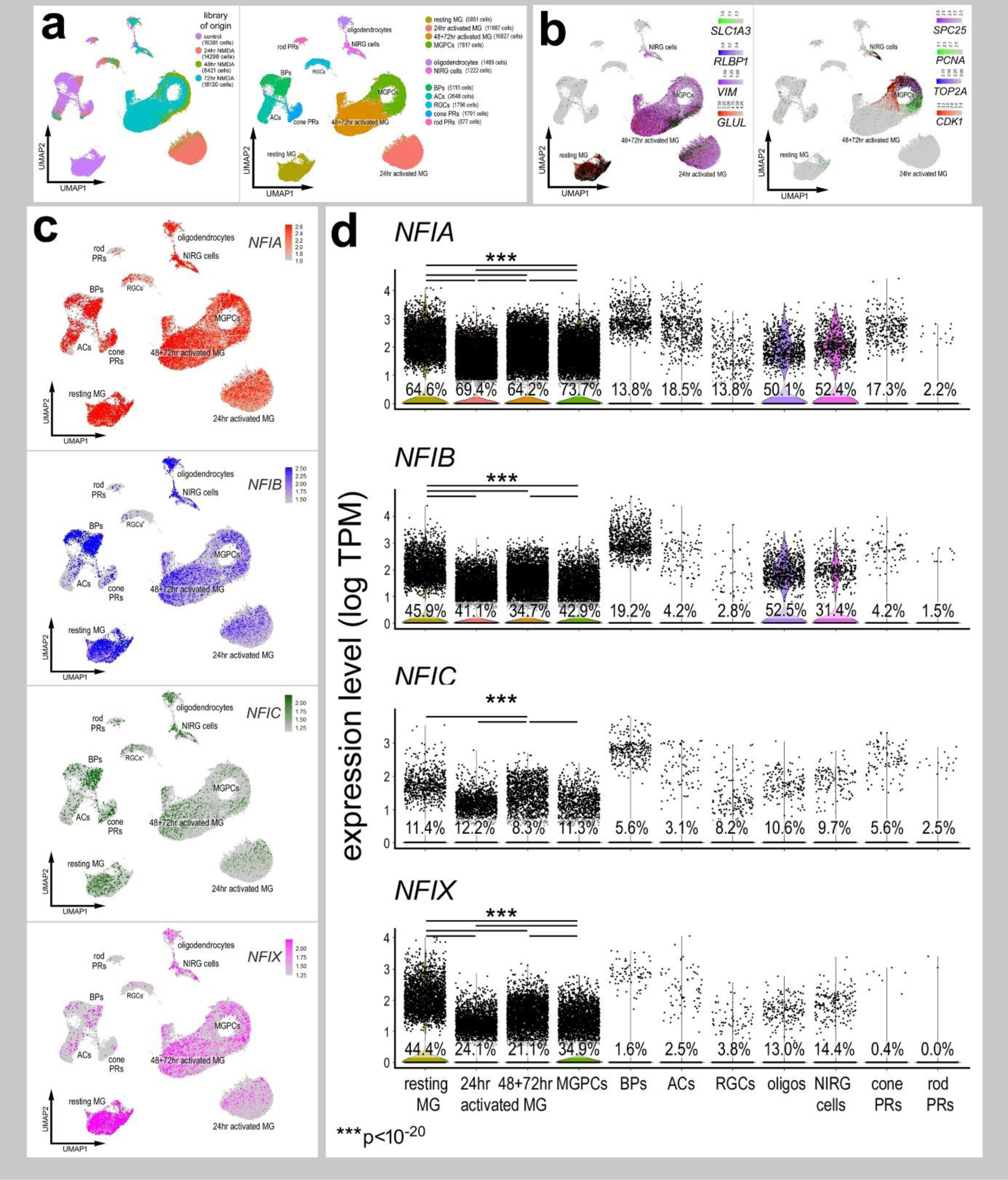
Patterns of expression of *NFIA, NFIB, NFIC* and *NFIX* in normal and NMDA-damaged chick retinas. scRNA-seq was used to identify patterns of expression of NFI’s among retinal cells with the data presented in UMAP (**a**-**c**) or violin plots (**d**). scRNA-seq libraries were aggregated for retinal cells from control eyes and eyes 24hr, 48hr, and 72hr after NMDA-treatment (**a**). UMAP-ordered cells formed distinct clusters of neuronal cells, resting MG, early activated MG, activated MG and MGPCs (**a,b**). Resting MG were identified based on elevated expression of *SLC1A3, RLBP1, VIM* and *GLUL* (**b**). Activated MG down-regulated *SLC1A3, RLBP1* and *GLUL*, and MGPCs up- regulated proliferation markers including *SPC25, PCNA, TOP2A* and *CDK1* (**b**). UMAP heatmaps of *NFIA, NFIB, NFIC* and *NFIX* demonstrate patterns of expression across retinal cell types. Violin plots illustrate the percentage of expressing cells, levels of gene expression and significant changes (***p<10exp-20) in levels (log TPM) were determined by using a Wilcox rank sum with Bonferoni correction.

*NFIA, NFIB, NFIC* and *NFIX* were not expressed a significant levels in retinal progenitor cells at E5 (Fig. 1c,d). Expression of *NFIC* and *NFIX* were low across all cell types during retinal development (Fig. 1c,d). By comparison, *NFIA* and *NFIB* were expressed by relatively few E8 progenitors, and at increasing levels and numbers of immature MG at E8 and maturing MG at E12 and E15 (Fig. 1c,d). In addition, *NFIA* and *NFIB* were detected in a few differentiating ganglion, amacrine, bipolar and photoreceptor cells at later stages (E12 and E15) of development (Fig. 1c).

To verify findings from scRNA-seq we applied antibodies to NFIA to retinal sections. Consistent with scRNA-seq data, NFIA-immunolabeling was absent from E5 retinas, but appeared at E6 in putative RPCs in the middle of prospective INL and differentiating neurons in the prospective IPL and GCL (Fig. 1e). At E8, NFIA-immunolabeling was present in the nuclei in putative RPCs, differentiating amacrine cells in the proximal INL, and differentiating ganglion cells (Fig. 1e). This pattern of labeling was similar at E10, with elevated levels seen in central retinal, similar levels in mid-peripheral retina, and very low levels in far peripheral retina (Fig. 1e). At E12 and E14, NFIA-immunofluorescence was present in the nuclei of scattered cells in the GCL and NFL, many presumptive amacrine cells in the proximal half of the INL, many presumptive differentiating MG in the middle of the INL, and scattered cells in the distal INL that likely were differentiating bipolar cells (Fig. 1e). By E16, patterns of immunolabeling for NFIA resembled that seen in post-hatch retina, with the exception of an absence of labeling in the ONL (Fig. 1e). According to scRNA-seq data, *NFIB* was detected in relatively few bipolar cells, rod photoreceptors and MG (Figs. 1c,d). Thus, we did not apply NFIB-antibodies to sections of embryonic retinas. Likewise, *NFIC* and *NFIX* were detected in very few embryonic retinal cells, and we thus did not probe for immunoreactivity.

### NFI’s in mature and damaged retinas

We next analyzed patterns of expression of NFIs in post-hatch chick retinas, wherein the cell types are mature and functional. We analyzed aggregate scRNA-seq libraries of cells from control and NMDA-damaged retinas at various time points (24, 48 and 72 hrs) after treatment (Fig. 2a). UMAP plots revealed clusters of cells that were identified based on well-established patterns of expression (Fig. 2a and Supplemental Fig. 2c). Resting MG formed a discrete cluster of cells and expressed high levels of *GLUL, RLBP1* and *SLC1A3* (Fig. 2b). After damage, MG down-regulate markers of mature glia as they transition into reactive glial cells and into progenitor-like cells that up-regulate *SPC25, PCNA, TOP2A* and *CDK1* (Fig. 2b).

Levels of *NFIA, NFIB, NFIC* and *NFIX* were relatively high in resting MG (Fig. 2c,d). Following NMDA-induced damage, the levels of all *NFIs* were reduced in MG and MGPCs at 24hrs, 48hrs, and 72hrs after treatment (Fig. 2c,d). The expression of *NFIA* and *NFIB* was prevalent in oligodendrocytes and Non-astrocytic Inner Retinal Glia (NIRGs) (Fig. 2c,d). NIRG cells are a distinct type of glial cell that has been described in the retinas of birds (Fischer et al., 2010a; Rompani and Cepko, 2010) and some types of reptiles (Todd et al., 2016b). *NFIC* and *NFIX* were expressed by relatively few oligodendrocytes and NIRG cells (Fig. 2d). In addition, we detected relatively high levels of *NFIA* and *NFIB* in some types of bipolar cells (Fig. 2c,d).

### Immunolabeling for NFIA and NFIB

To validate patterns of expression for NFIA and NFIB we performed immunofluorescence labeling in post-hatch chick retina. Immunofluorescence for NFIA was observed in different types of retinal neurons and glia. NFIA was detected in many AP2α-positive amacrine cells (Fig. 3a) and a few Brn3a-positive ganglion cells (Fig. 3b). Further, NFIA was detected in bipolar cells in the distal INL that were negative for Islet1, but positive for Otx2 (Fig. 3c,d). All Sox2-positive MG were co-labeled for NFIA, whereas cholinergic amacrine cells known to express Sox2 and Islet1 (Stanke et al., 2008), were negative for NFIA (Fig. 3e). In addition, many Nkx2.2-positive NIRG cells in the IPL were co-labeled for NFIA (Fig. 3f).

**Figure 3.**
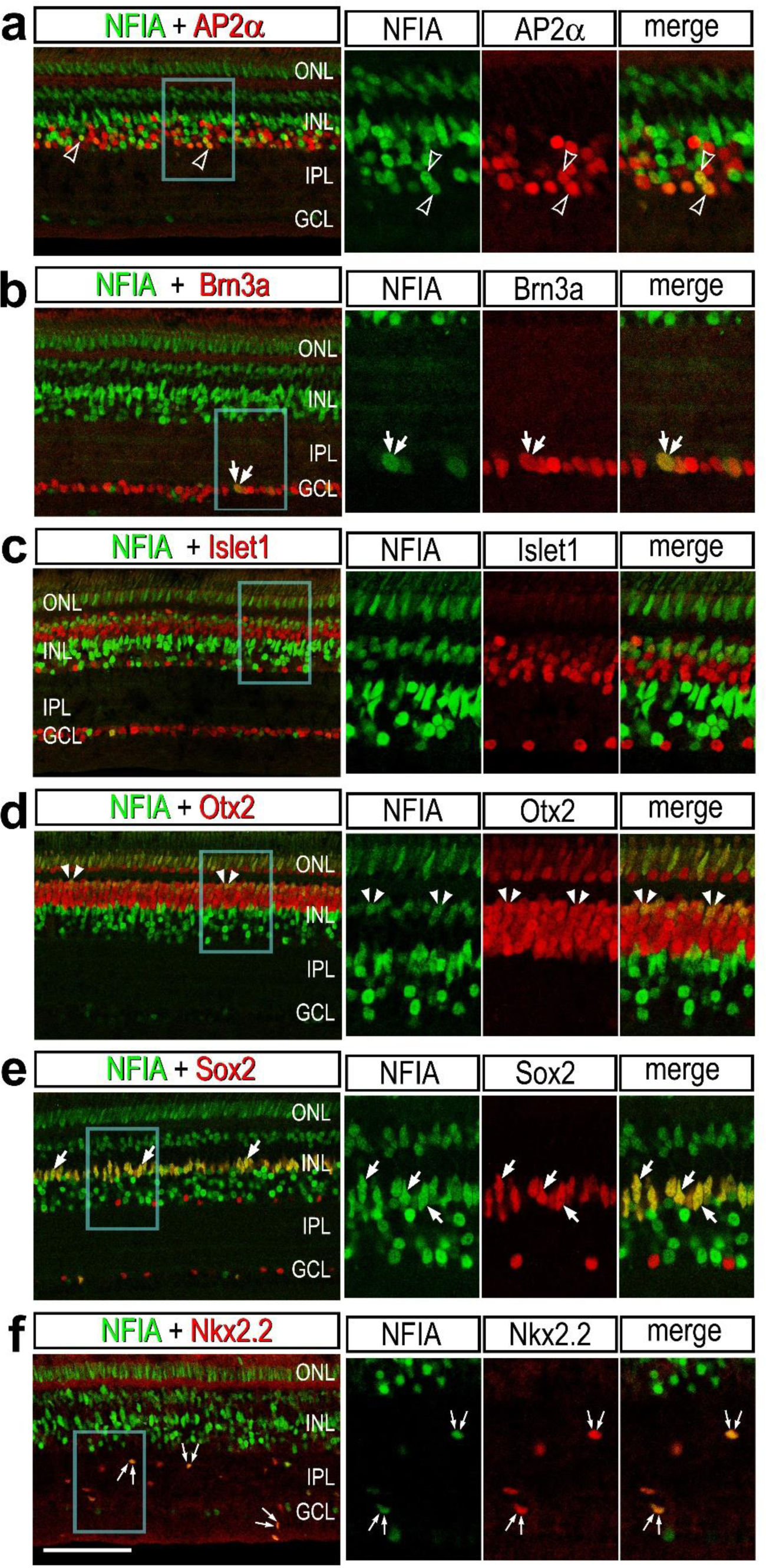
Immunofluorescence for NFIA in different types of cells in chick retina. Retinas were obtained from eyes that were injected with saline (**a**-**e**) or NMDA (**f**) and harvested 48 hrs after treatment. Vertical sections of the retina were labeled with antibodies to NFIA (green) and AP2α (**a**), Brn3a (**b**), Islet1 (**c**), Otx2 (**d**), Sox2 (**e**) or Nkx2.2 (**f**; red). Arrows indicate the nuclei of MG, small double arrows indicate the nuclei of NIRG cells, arrow heads indicate the nuclei of microglia, hollow arrow heads indicate nuclei of amacrine cells, small double-arrow heads indicate nuclei of bipolar cells, and double arrows indicate nuclei of ganglion cells. Areas indicated by blue boxes are enlarged 2-fold in the panels to the right. The calibration bar (50 µm) applies to all panels. Abbreviations: ONL – outer nuclear layer, INL – inner nuclear layer, IPL – inner plexiform layer, GCL – ganglion cell layer.

Immunolabeling for NFIB revealed patterns of expression that closely matched those for scRNA-seq. NFIB-immunoreactivity was detected in; (i) a subset of amacrine cells that were Ap2α-positive, (ii) bipolar cells that were Otx2-positive and Islet1- negative), and (iii) all MG that were Sox2-positive (Fig. 4a,b). To determine whether NFIB-immunoreactivity was present in microglia and NIRG cells, we labeled NMDA- damaged retinas. All CD45+ microglia and Nkx2.2+ NIRG cells in the inner retina were labeled for NFIB at 24hrs after NMDA-treatment (Fig. 4b)

**Figure 4.**
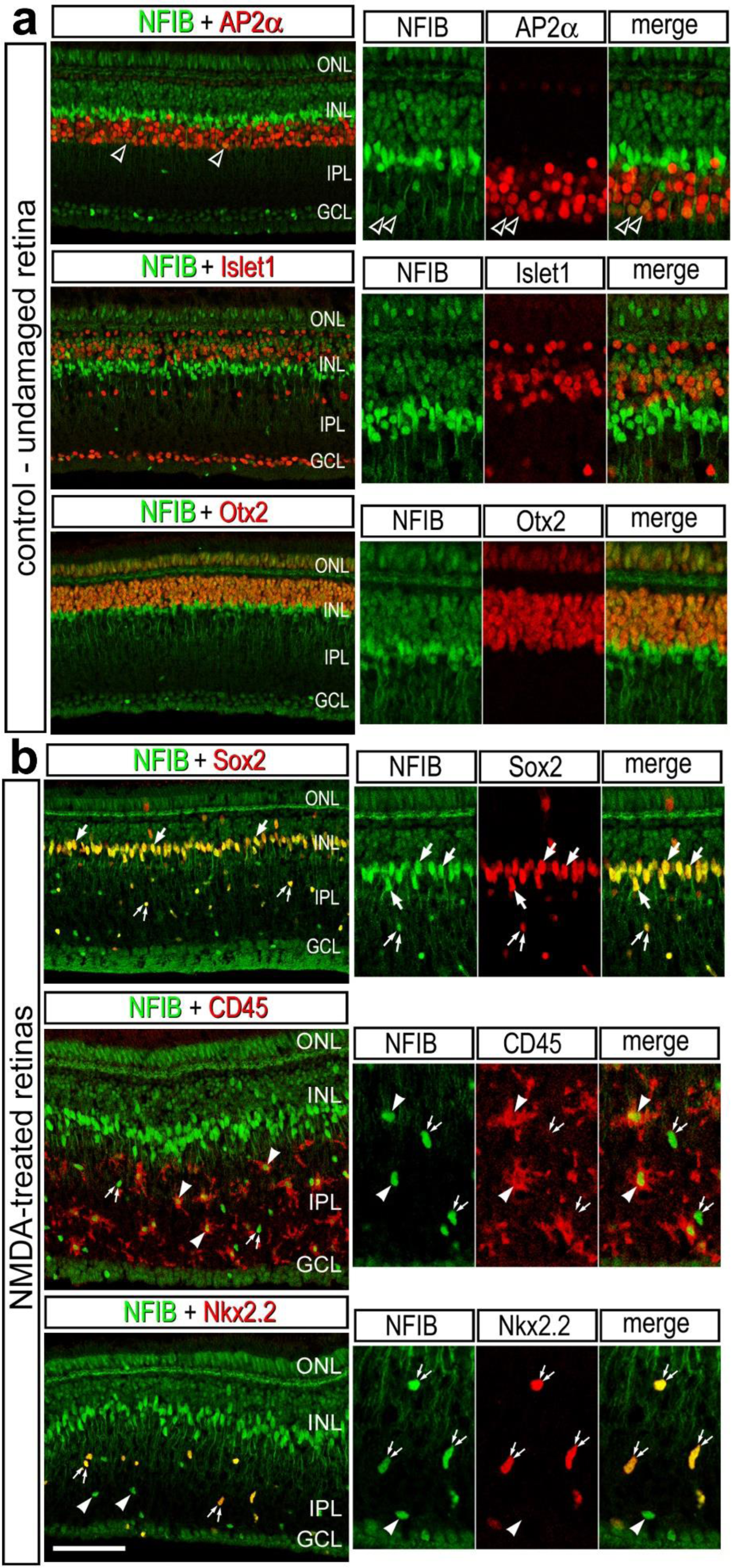
Immunofluorescence for NFIB in different types of cells in chick retina. Retinas were obtained for eyes injected with saline (**a**) or NMDA (**b**) at 48hrs after treatment. Vertical sections of the retina were labeled with antibodies to NFIB (green) and AP2α (**a**), Islet1 (**a**), Otx2 (**a**), Sox2 (**b**), CD45 (**b**) or Nkx2.2 (**b**; red). Arrows indicate the nuclei of MG, small double arrows indicate the nuclei of NIRG cells, arrow heads indicate the nuclei of microglia, and hollow arrow heads indicate nuclei of amacrine cells. Panels to the right are enlarged 2-fold from the panels to the left. The calibration bar (50 µm) applies to all panels to the left. Abbreviations: ONL – outer nuclear layer, INL – inner nuclear layer, IPL – inner plexiform layer, GCL – ganglion cell layer.

To further discriminate NFIA and NFIB in different types of bipolar cells we re-embedded scRNA-seq data for bipolar cells from control retinas. Unsupervised UMAP- ordering of cells revealed 11 distinct clusters of bipolar cells, based on patterns of expression for genes such as *CALB2, ISL1, ARR3, SPON1, ALCAM, BHBHE23, PROX1* and *GRIK1* (Fig. 5a-d). Consistent with recent findings from embryonic chick retinas (Yamagata et al., 2021) all clusters of bipolar cells were positive for *VSX1, VSX2* and *OTX2*, and a binary segregation of types based on expression of *ISL1* or *GRIK1* (ON vs OFF bipolar cells). Clusters of cells that expressed *NFIA* overlapped with those that expressed *NFIB*, with the exception of cluster 8. *NFIA* and *NFIB* were predominantly expressed in clusters 4, 5, 6 and 11, which overlapped with expression of *GRIK1* (Glutamate Ionotropic Receptor Kainate Type Subunit 1; Fig. 5c-f, suggesting that these cells were OFF-center bipolars. By comparison Yamagata and colleagues (Yamagata et al., 2021) described 22 distinct types of bipolar cells in E16 chick retinas. Collectively, the findings from immunolabeling studies correlated very closely with the patterns of expression reveal by scRNA-seq. Immuolabeling for NFIA and NFIB in NMDA-damaged retinas failed to reveal significant differences in immunofluorescence in the nuclei of MG (not shown).

**Figure 5.**
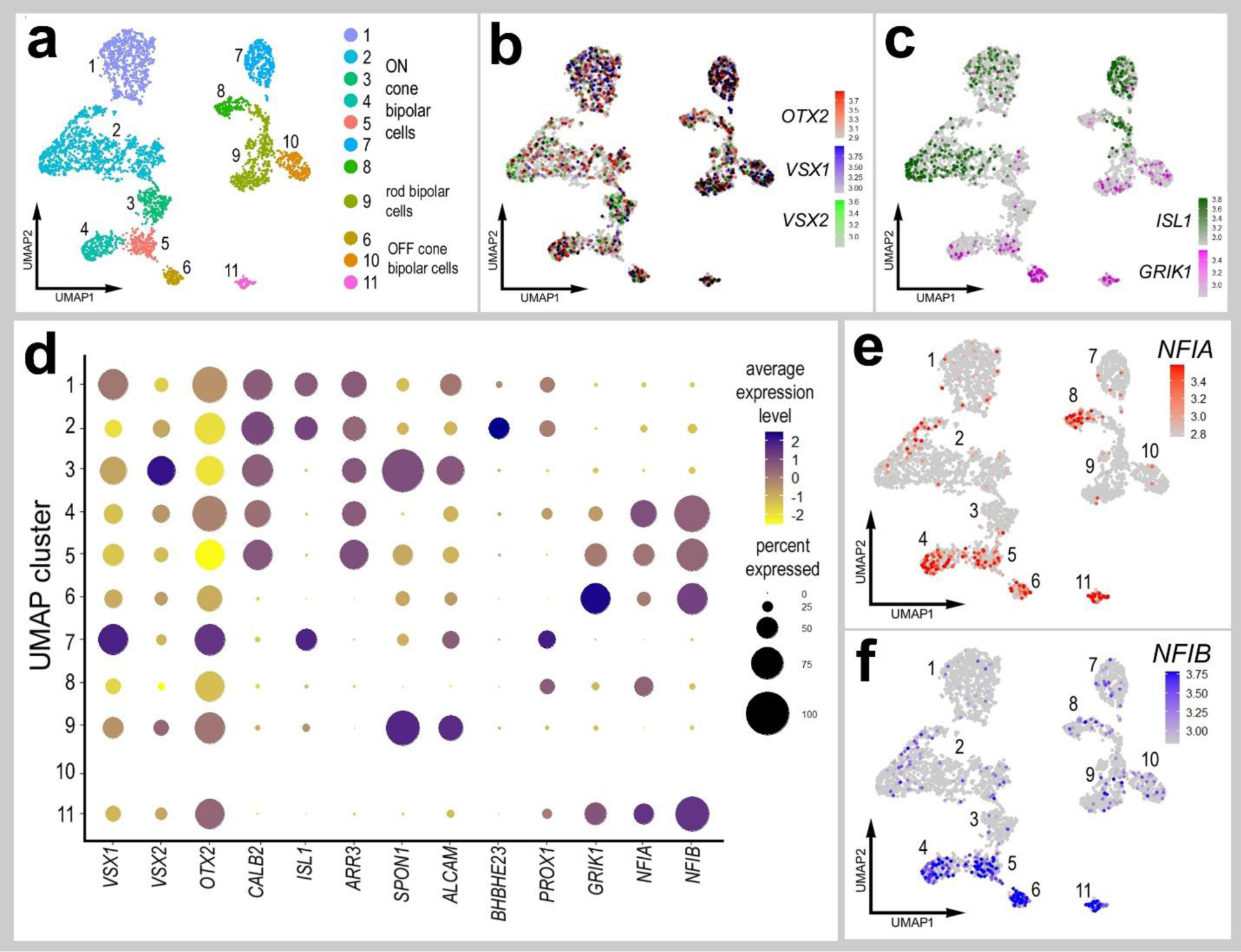
Expression levels of NFIA and NFIB in different types of bipolar cells. Bipolar cells were isolated from aggregate scRNA-seq libraries from saline-treated retinas and re-embedded in UMAP plots. UMAP ordering of bipolar cells revealed 11 distinct clusters of cells (**a**). All bipolar cell clusters contained cells that expressed *OTX2, VSX1* and *VSX2* (**b**). ON- and OFF-center bipolar cells were distinctly clustered based on expression of *ISL1* or *GRIK1* (**c**). The dot plot in **d** illustrates the average expression level (heatmap) and percent expression (dot diameter) for bipolar ell markers that distinguished the different clusters of cells. The expression of *NFIA* and *NFIB* is illustrated UMAP heatmap and dot plots in distinct clusters of bipolar cells (**d**-**f**).

### NFI’s in eRPCs, MG, MGPCs and CMZ progenitors

We next isolated MG, aggregated and normalized scRNA-seq data for cells from saline-, NMDA-, FGF2+insulin- and NMDA+FGF2+insulin-treated retinas to directly compare levels of NFIs. UMAP plots revealed distinct clustering of MG from control retinas and MG from 24hrs after NMDA-treatment, whereas MG from retinas at 48 and 72hrs after NMDA and from retinas treated with insulin and FGF2 formed a large cluster with distinct regions (Fig. 6a-e). UMAP plots revealed distinct patterns of expression of genes associated with resting MG, de-differentiating MG, activated MG and MGPCs (Fig. 6c-e and Supplemental Fig. 2d). Different zones, representing MGPCs in different phases of the cell cycle were comprised of cells from different times after NMDA- treatment and FGF2+insulin-treatment (Fig. 6b,e). Expression of *NFIs* was most widespread and elevated in MG in undamaged retinas compared to MG from retinas treated with NMDA and/or insulin and FGF2 (Fig. 6f). Levels of NFIs were down- regulated in MG by treatment with NMDA and/or insulin+FGF2, with the largest decreases in prevalence and levels seen with insulin+FGF2-treatment and in the MGPC3 cluster of cells (Fig. 6f).

**Figure 6.**
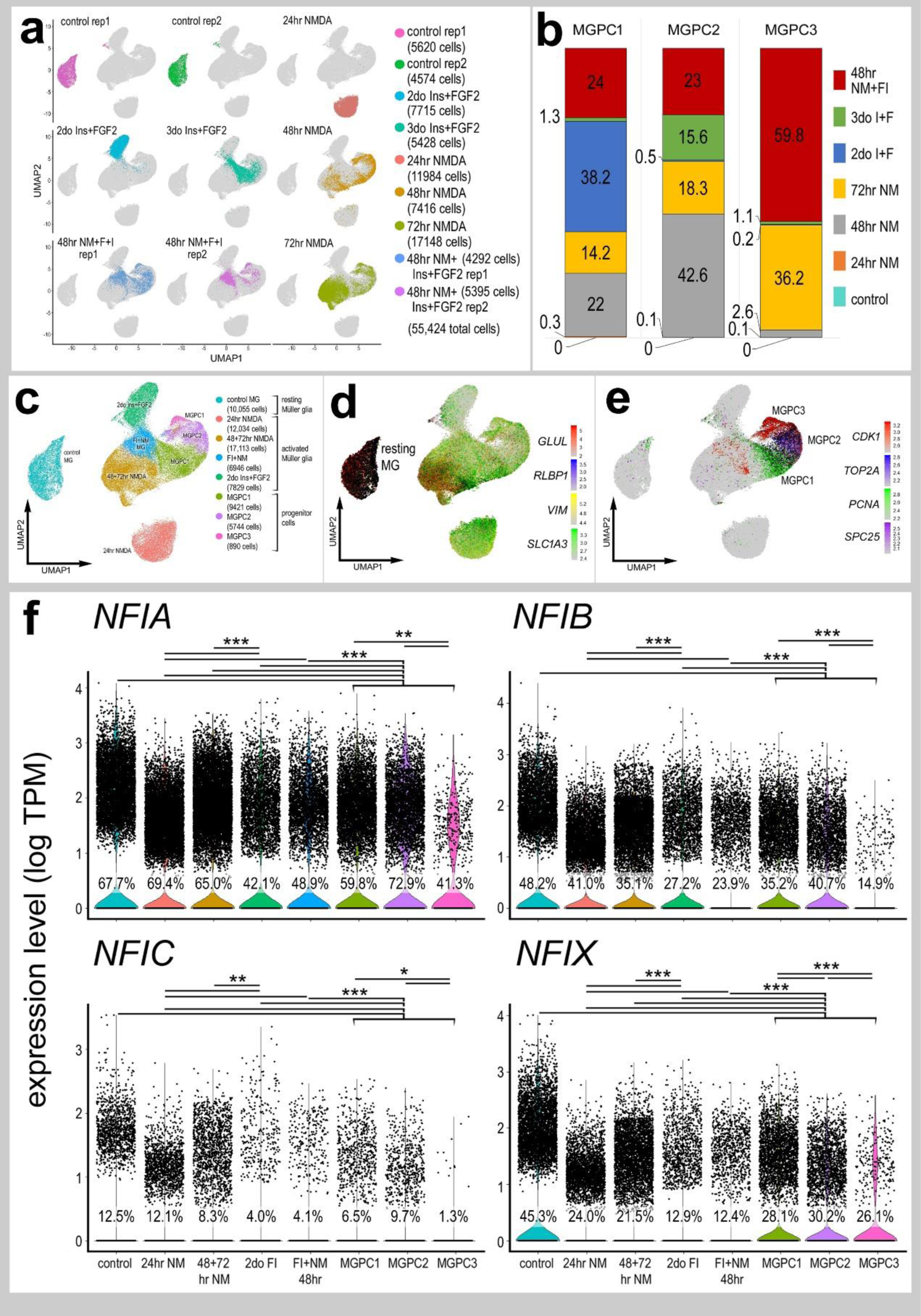
Comparison of NFI expression levels in Müller glia and MGPCs from different treatment conditions. scRNA-seq was used to identify patterns of expression of NFIs in MG and MGPCs at different time points after NMDA damage and/or FGF + insulin growth factor treatment. Nine different libraries were aggregated for a total of more than 55,000 MG and MGPCs (**a,c**). MGPCs were identified based on high-levels of expression of *CDK1, TOP2A, PCNA* and *SPC25* (**c**,**e**), and were a mix of cells from 48hrs NMDA+FGF2+insulin, 48hrs NMDA, 72hrs NMDA and 3doses of insulin+FGF2 (**b**). Resting MG were identified based on expression of *GLUL, RLBP1, SLC1A3* and *VIM* (**c,d**). Each dot represents one cell and black dots indicate cells with 2 or more genes expressed. The expression of *NFIA, NFIB, NFIC* and *NFIX* is illustrated violin plots with population percentages and statistical comparisons (**f**). Significance of difference (*p<0.01, **p<10^-10^, ***p<10^-20^) in expression levels (log TPM) were determined by using a Wilcox rank sum with Bonferoni correction. Abbreviations: ONL – outer nuclear layer, INL – inner nuclear layer, IPL – inner plexiform layer, GCL – ganglion cell layer.

To directly compare levels of NFIs in MGPCs and embryonic retinal progenitor cells (eRPCs) we isolated, aggregated and normalized scRNA-seq data for these cells. UMAP clustering of cells revealed distinct separation of proliferating embryonic RPCs and MGPCs (Fig. 7a-c). Although *NFIA* and *NFIB* were detected in E8 RPCs at high levels, the prevalence of expression among MGPCs was significantly higher (Fig. 7d,f). Levels of *NFIC* and *NFIX* were negligible in early and late RPCs, whereas levels were elevated and more wide-spread in MGPCs (Fig. 5(7e,f).

**Figure 7.**
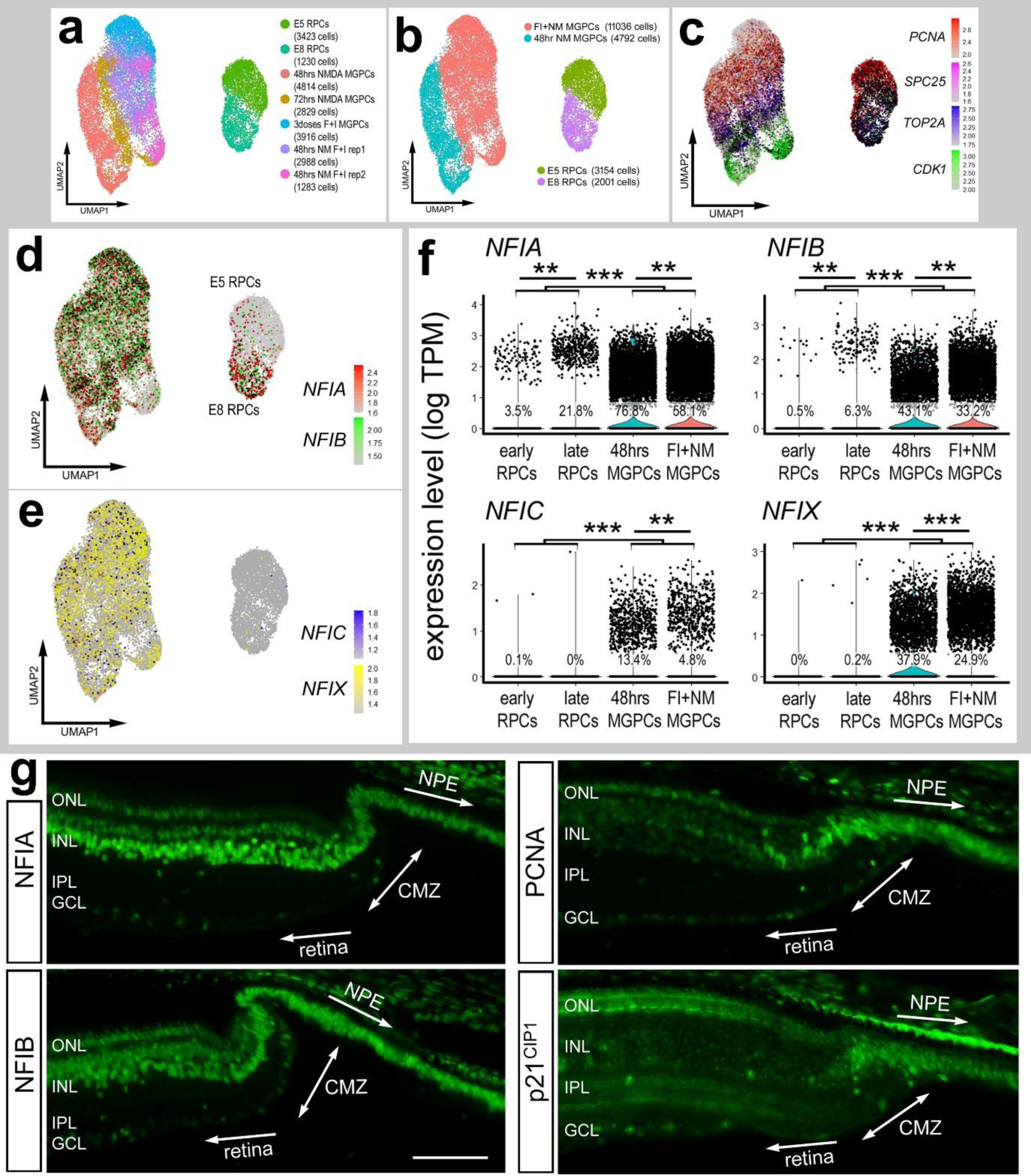
Expression levels of NFI’s in embryonic RPCs, Müller glia, MGPCs and CMZ progenitors. scRNA-seq was used to compare patterns of expression of NFIs in embryonic retinal progenitor cells (eRPCs) and MGPCs from different time points after NMDA-damage and/or treatment with insulin+FGF2 (**a**). Seven different scRNA-seq libraries were aggregated for a total of 20,483 eRPCs and MGPCs (**a,b**). eRPCs and MGPCs expressed high levels of proliferation-associated genes including *PCNA, SPC25, TOP2A* and *CDK1* (**c**). The expression of *NFIA, NFIB, NFIC* and *NFIX* is illustrated in UMAP heatmap plots and violin plots with population percentages and statistical comparisons (**d**-**f**). Significance of difference (**p<10exp-10, ***p<10exp-20) in expression levels (log TPM) were determined by using a Wilcox rank sum with Bonferoni correction. **g** – Sections of the far peripheral retina were label with antibodies to NFIA, NFIB, PCNA and p21cip1. The calibration bar represents 50 µm. Abbreviations: ONL – outer nuclear layer, INL – inner nuclear layer, IPL – inner plexiform layer, GCL – ganglion cell layer, CMZ – circumferential marginal zone, NPE – non-pigmented epithelium.

We next examined whether retinal progenitors in the circumferential marginal zone (CMZ) expressed NFIA and NFIB. The eyes of hatched chicks are known to contain a zone of proliferating neuronal progenitors at the far peripheral edge of the retina (Fischer and Reh, 2000; Fischer et al., 2002a). These progenitors proliferate slowly, but can be stimulated by insulin and FG2 to proliferate at increased rates and generate inner retinal neurons and ganglion cells (Fischer and Reh, 2000; Fischer et al., 2002a). Our scRNA-seq databases did not include CMZ progenitors at the far peripheral edge of the retina. Thus, we immunolabeled sections of CMZ with antibodies to NFIA and NFIB. Immunofluorescence for NFIA and NFIB was detected in the nuclei of cells in the non-pigmented epithelium (NPE) of the ciliary body (Fig. 7g). In addition nuclear labeling for NFIA and NFIB was observed in cells through the CMZ and in numerous cells in the INL and ONL of the neural retina (Fig. 7g). The CMZ cells were identified by labeling for PCNA and p21^cip1^ (Fig. 7g).

### NFIs in pig and monkey retinas

We next sought to characterize the patterns of expression of NFIs in the retinas of large mammals. We analyzed retinal sections and scRNA-seq databases for adult pig and adult macaque monkeys. We aggregated scRNA-seq libraries from 4 normal adult pig retinas for a total of 15,236 cells (Fig 8a), after filtering out doublets, cells with low genes per cell and elevated mitochondrial RNA content. Different retinal cell types were ordered into distinct UMAP clusters, including 2 clusters of MG (Fig. 8b). The clusters of MG were segregated into resting and activated, with the activated cells expressing elevated levels of *ATF3*, *FOSB, EGR1, TCF7, FGFR1* and *PAK3* (Supplemental Fig. 3). We speculated that activated MG resulted from the process of tissue harvesting and tissue dissociation prior to capture with scRNA-seq reagents. We found that *NFIA* and *NFIB* were predominantly expressed in MG, but were also expressed by a few amacrine and bipolar cells (Fig. 8c). By comparison, *NFIC* and *NFIX* had relatively wide-spread expression in rod and cone photoreceptors, microglia, amacrine cells and bipolar cells (Fig. 8c). In addition, *NFIC* and *NFIX* were detected in resting and activated MG (Fig. 8c).

**Figure 8.**
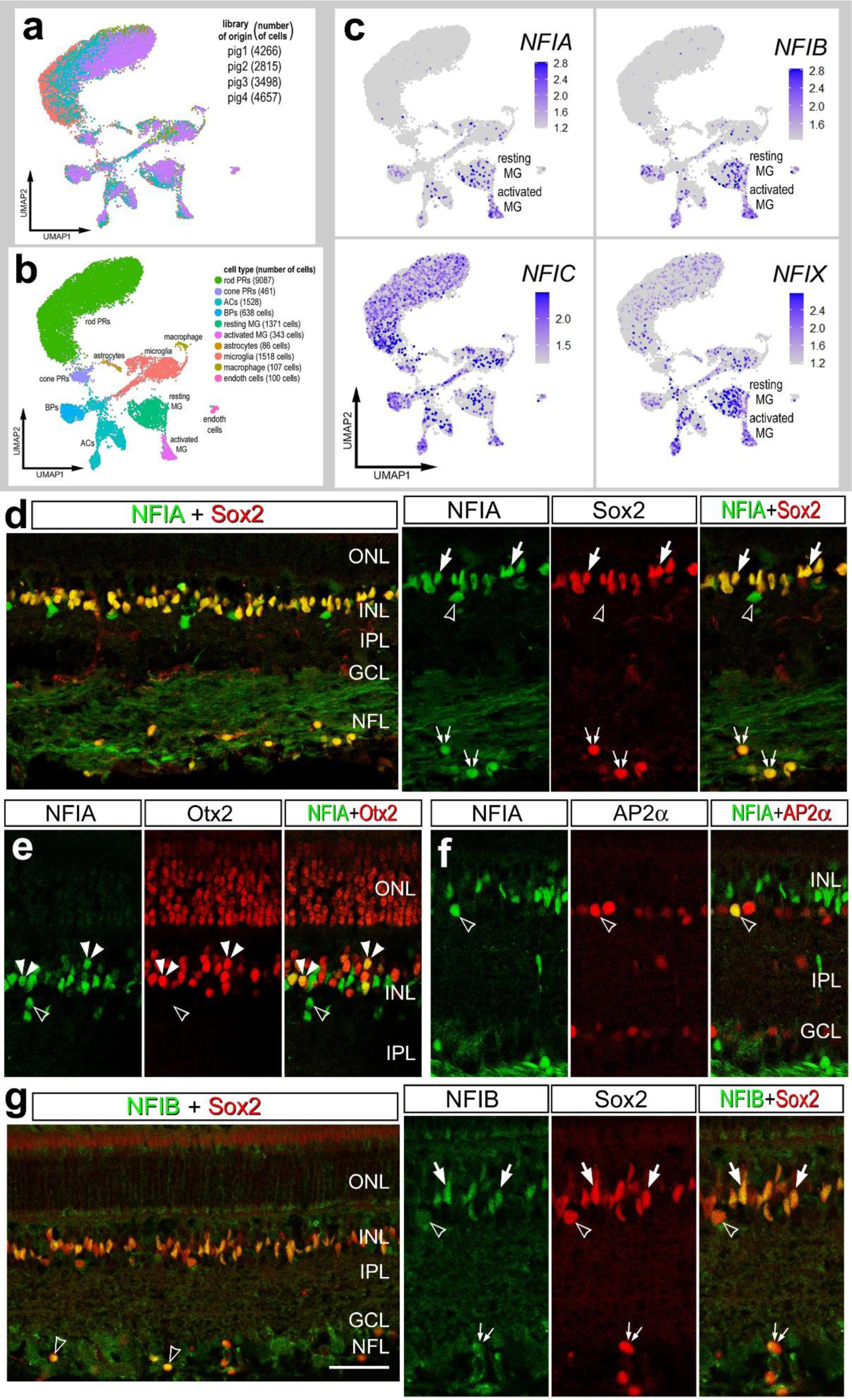
NFI’s in the pig retina. scRNA-seq was used to identify patterns of expression of NFIs in adult pig retina. Four different scRNA-seq libraries were aggregated for a total of 15,236 cells (**a**). UMAP clustered cells were identified based on cell-distinguishing markers (**b**). The expression of *NFIA, NFIB, NFIC* and *NFIX* is illustrated in UMAP heatmap plots (**c**). Vertical sections of central and peripheral regions of the retina were labeled with antibodies to NFIA (green; **d**-**f**), NFIB (green; **g**), Sox2 (red; **d** and **g**), Otx2 (red; **e**), or AP2α (red; **f**). Arrows indicate the nuclei of MG, small double arrows indicate the nuclei of an unidentified cell type in the NFL, hollow arrow heads indicate nuclei of amacrine cells, and small double-arrow heads indicate nuclei of bipolar cells. The calibration bar represents 50 µm. Abbreviations: ONL – outer nuclear layer, INL – inner nuclear layer, IPL – inner plexiform layer, GCL – ganglion cell layer.

Consistent with scRNA-seq data from pig retinas, we detected immunofluorescence for NFIA in the nuclei of all Sox2-positive MG in the INL and in unidentified cells in the NFL (Fig. 8d). In addition, NFIA was detected in Otx2-positive bipolar cells, but only in peripheral regions of the retina (Fig. 8e). NFIA was detected in a few sparsely distributed cells in the amacrine cell layer of the INL and some of these cells were co-labeled for AP2α (Fig. 8f). By comparison, NFIB-immunofluorescence was detected in the nuclei of all Sox2+ MG in the INL and in sparsely distributed cells in the NFL (Fig. 8g). In addition, NFIB was detected in Sox9-positive nuclei that were scattered in the NFL (Fig. 8g). The identity of the NFIB/NFIA/Sox2-positive cells in the NFL remains uncertain.

We next characterized the expression patterns of NFIs in the retina of the macaque monkey. For the monkey we aggregated scRNA-seq libraries from 2 normal adult retinas for a total of 14,287 cells after filtering out doublets, cells with low genes per cell and elevated mitochondrial RNA content. Different retinal cell types were ordered into distinct UMAP clusters based on patterns of expression of cell-identifying markers such as *GNB3, PAX6, GRIK1* and *RLBP1* (Fig. 9a,b). Similar to the patterns seen in the chick retina, *NFIA* and *NFIB* were detected in MG, rod bipolar cells, OFF bipolar cells and amacrine cells (Fig. 9c,d). *NFIC* and *NFIX* were detected in MG, rod bipolar cells, amacrine cells and OFF bipolar cells (Fig. 9e,f). The expression of *NFIC* was also prominent in rod photoreceptors (Fig. 9e).

**Figure 9.**
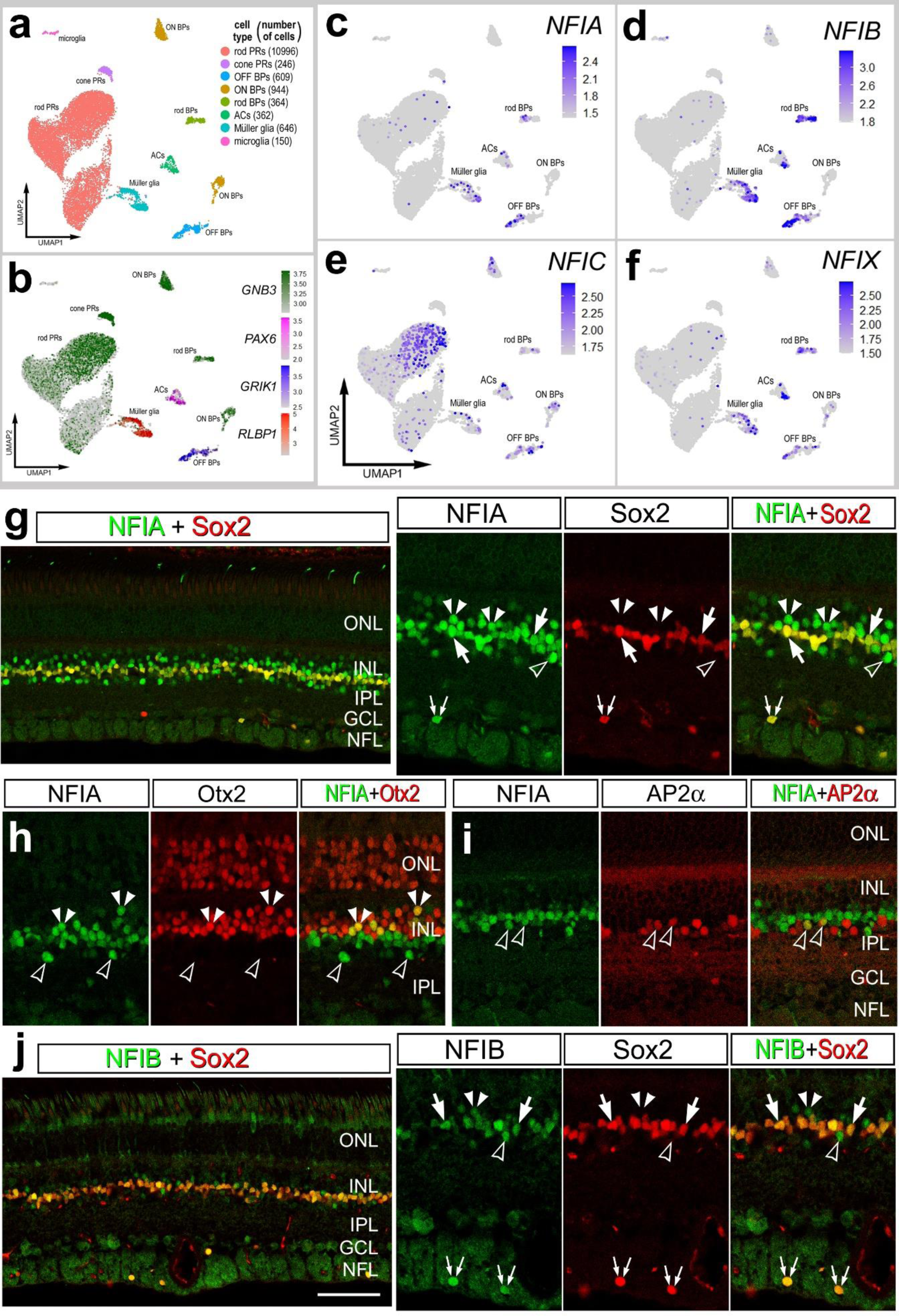
NFIs in the monkey retina. scRNA-seq was used to identify patterns of expression of NFIs in adult monkey retina. Two different scRNA-seq libraries were aggregated for a total of 14,319 cells (**a**). UMAP clustered cells were identified based on cell-distinguishing markers (**b**). The expression of *NFIA, NFIB, NFIC* and *NFIX* is illustrated in UMAP heatmap plots (**c-f**). Vertical sections of central and peripheral regions of the retina were labeled with antibodies to NFIA (green; **g**-**i**), NFIB (green; **j**), Sox2 (red; **g** and **j**), Otx2 (red; **h**), or AP2α (red; **i**). Arrows indicate the nuclei of MG, small double arrows indicate the nuclei of an unidentified cell type in the NFL, hollow arrow heads indicate nuclei of amacrine cells, and small double-arrow heads indicate nuclei of bipolar cells. e. The calibration bar represents 50 µm. Abbreviations: ONL – outer nuclear layer, INL – inner nuclear layer, IPL – inner plexiform layer, GCL – ganglion cell layer.

Consistent with scRNA-seq data from the monkey retina, NFIA-immunolabeling was detected in the nuclei of all Sox2-positive MG (Fig. 9g). In addition, NFIA was detected in a many cells in amacrine and bipolar cell layers of the INL (Fig. 9g-i). We observed sparsely distributed NFIA+ cells in the NFL that were co-labeled for Sox2(Fig. 9d). Many of the NFIA+ cells in the distal INL were bipolar cells that were co-labeled for Otx2 (Fig. 9h), and many of the NFIA+ cells in the proximal INL were amacrine cells that were co-labeled for AP2α (Fig. 9i). NFIB-immunofluorescence was detected in the nuclei of all Sox2+ MG in the INL and sparsely distributed Sox2+ cells in the NFL (Fig. 9j). In addition, relatively low levels of NFIB were detected in presumptive amacrine and bipolar cells in the INL (Fig. 9j). The identity of the NFIA/NFIB/Sox2-positive cells in the NFL remains uncertain.

## Discussion

This study demonstrates that NFI transcription factors are expressed by MG and by developing and differentiated neurons in the retinas of chicks, pigs and non-human primates. NFI expression is decreased in proliferating MGPCs in chick retinas. These observations are consistent with a role for NFI factors in promoting or maintaining differentiated neuronal or glial phenotype and limiting progenitor cell competence. NFIs are not likely to promote differentiated cells by forcing cells to exit the cell cycle, as these factors are widely expressed by proliferating MGPCs. This is in agreement with results from mouse retina, where overexpression of NFI gene products resulted in increases in numbers of MG and bipolar cells and deletion of NFI genes resulted in an increase in late progenitors at the expense of late-born retinal cell types (Clark et al., 2019).

NFIs may not suppress proliferation, but instead may suppress neurogenic potential or limit potential to late-born retinal cell types. NFIs are highly expressed by resting MG, but levels are reduced while remaining wide-spread in activated MG, prior to re-entering the cell cycle and by proliferating MGPCs. Similarly, proliferating CMZ retinal progenitors expressed NFIA and NFIB, and these progenitors are known to produce late-born retinal cell types such as amacrine cells and MG (Fischer and Reh, 2000). We observed transient decreases in expression levels for all NFIs from resting MG to activated MG at 24hrs after NMDA-treatment. This observation is consistent with the hypothesis that de-differentiation occurs during activation of MG and transition to progenitor-like phenotype. We propose that transient down-regulation of NFIs is part of the process of MGPC formation. The largest decreases in numbers of MG that expressed NFIs was observed in retinas treated with insulin and FGF2. The combination of insulin and FGF2 is known to stimulate the formation of proliferating MGPCs in the absence of retinal damage (Fischer et al., 2002b, 2009). FGF2 is potently neuroprotective while stimulating the formation of MGPCs when applied before NMDA- induced damage (Fischer et al., 2014). Further, FGF2 is known to suppress the expression of glial genes during late stages of embryonic development (Kruchkova et al., 2001). Similarly, our data from scRNA-seq indicate significant down-regulation of mRNA’s for mature glial markers, such as glutamine synthetase, GLAST1, and RLBP1 in MG treated with 2 doses of insulin and FGF2 without reprogramming into proliferating MGPCs (Campbell et al., 2019, 2021; Hoang et al., 2020; Palazzo et al., 2020).

Collectively, the down-regulation of NFI’s in MG treated with insulin and FGF2 is consistent with the hypothesis that these growth factors drive the process of de- differentiation and this process involves reducing levels of NFIs.

### Comparison of expression patterns of NFIA and NFIB in the retinal neurons in chick, pig and monkey retinas

Patterns of expression of NFI were similar across cell types in the retinas of chicks, pigs and monkeys. These patterns of expression were similar to those reported for cells in the mouse retina (Keeley and Reese, 2018; Clark et al., 2019). For example, NFIA and NFIB were prominently expressed in MG in all species. However, there were some notable exceptions in differences in patterns of NFIs. NFIA was detected in cone photoreceptors in chick, but was not detected in photoreceptors in pig and monkey retinas. Microglia in the chick retina were immunoreactive for NFIA and NFIB, whereas NFIC and NFIX were detected in pig microglia and NFI’s were not detected in the microglia from the monkey retina. Expression of NFIA in bipolar and amacrine cells was prevalent in chick and monkey retinas, but not in pig retinas. The identity of the NFIA/NFIB/Sox2/Sox9-positive cells in the NFL remains uncertain. It is likely that these cells are astrocytes. In the retinas of mice, dogs and monkeys, astrocytes scattered in the NFL are known to be immunoreactive for Sox2, Sox9, S100β, Pax2 and GFAP (Fischer et al., 2010b; Stanke et al., 2010).

### NFIs in the chick CMZ

Expression of NFIA and NFIB in the NPE of the ciliary body immediately anterior to the CMZ and within the neural retina suggests that the functions of NFIs are pleiotropic and context dependent. It is possible that the functions of NFIs in the NPE includes suppression of neurogenic potential. The NPE is derived from the neural tube but neurogenesis and proliferation is suppressed during development. However, treatment with different growth factors can stimulate proliferation and *de novo* neurogenesis from NPE cells *in vivo* in the chick model (Fischer and Reh, 2003). In addition, NFIA and NFIB were detected in CMZ progenitors. These progenitors proliferate at relatively low levels and produce MG and inner retinal neurons, but can be stimulated by growth factor to proliferate at increased rates and generate retinal neurons (Fischer and Reh, 2002, 2003). It is possible that the expression of NFIs by CMZ progenitors acts to restrict competency to late-born cell types such as bipolar cells, amacrine cells and MG.

### NFIs in mature neurons and glia

The functional roles for NFIs has been well-studied during the development of the CNS. However, the functional roles of NFIs remains uncertain in adult neuronal tissues. During development combined conditional knock-out of *Nfia, Nfib* and *Nfix* in radial glial cells prevents neuronal and glial differentiation, whereas deletion of individual NFI’s tends to result in delayed glial and neuronal differentiation (reviewed by Harris et al., 2015). The influence of NFIs on glial cells is likely coordinated with other transcription factors and cell signaling pathways that influence glial specification. In cortical stem cells, for example, *Ezh2* a methyltransferase in the polycomb repressor complex 2 (PRC2) and *Sox9* are repressed by NFIB and NFIX (Heng et al., 2014). Ezh2 activity is required for the normal formation of MG in the retina (Iida et al., 2015) and Sox9, along with other SoxE factors, promote progenitor and glial identity in a context-specific manner (Weider and Wegner, 2017). By comparison, *Hes1* and *Hes5*, bHLH factors down-stream of Notch-signaling are repressed by *Nfia* and *Nfib* (Piper et al., 2010). Hes1 and Hes5 promote specification of MG in developing rodent retinas (Furukawa et al., 2000; Hojo et al., 2000; Ohtsuka et al., 2001). Similarly, NFIA has been shown to promote glial cell fate and favor astrocyte specification in the developing rodent brain (Deneen et al., 2006; Namihira et al., 2009; Kang et al., 2012; Glasgow et al., 2014).

Although the roles of NFIs in developing neural tissues have been well-established, the roles of NFIs in mature neurons and glia remains less certain. The conditional deletion of NFIA from mature astrocytes influences responses to injury (Laug et al., 2019) and the activity of neuronal circuits (Huang et al., 2020). We have recently reported that conditional deletion of *Nfia, Nfib* and *Nfix* from mouse MG permits the reprograming of these glia into neuron-like cells when retinas were damaged and treated with insulin and FGF2 (Hoang et al., 2020). In the chick retina, we observed elevated levels of *NFIA* and *NFIB* in mature MG and lower levels in MGPCs, consistent with the hypothesis that these factors limit the neurogenic potential of MGPCs. For example, NFIA has been shown to promote glial cell fate and favor astrocyte specification in the developing rodent brain (Deneen et al., 2006; Namihira et al., 2009; Kang et al., 2012; Glasgow et al., 2014). Other transcription factors that may limit neurogenic potential of neuronal progenitors can include Id1, Id4, Sox2, Sox8, Sox9, Sox10, Hes1 and Hes5 (Imayoshi et al., 2013; Weider and Wegner, 2017; Akdemir et al., 2020). These factors are also expressed by chick MGPCs (Palazzo et al., 2020; Campbell et al., 2021), suggesting redundancy and a network of transcription factors that act to maintain glial phenotype and suppress neuronal differentiation. Not surprisingly this network of pro-glial factors is not prominent in MGPCs in damaged zebrafish retinas (Hoang et al., 2020).

### Conclusions

NFIs are expressed by many different types of retinal neurons and glia during development and in mature retinas. NFI expression is maintained in some types of mature neurons and all types of glia, including oligodendrocytes, microglia and MG. Patterns of expression of NFIs are very similar in the retinas of chicks, pigs and monkeys. NFIA and NFIB are prominently expressed by MG in the retinas of all species, and likely at to maintain glial phenotype. In chick retinas, levels of NFIA and NFIB are down-regulated during activation and de-differentiation of MG and in proliferating MGPCs. Our findings are consistent with the notion that NFIs act to maintain phenotype in mature retina neurons and glia.

### Data availability

RNA-Seq data for gene-cell matrices are deposited in GitHub https://github.com/jiewwwang/Single-cell-retinal-regeneration https://github.com/fischerlab3140/scRNAseq_libraries scRNA-Seq data can be queried at https://proteinpaint.stjude.org/F/2019.retina.scRNA.html.

## Author Contributions

HME and WAC designed and executed experiments, prepared scRNA-seq libraries, performed bioinformatic analyses, gathered data and constructed figures. LEK and ECH executed experiments and gathered data. AJ and MAM provided pig retinas. MS and KM prepared scRNA-seq libraries for pig and monkey retinas.AJF designed experiments, analyzed data, performed bioinformatic analyses, constructed figures and wrote the manuscript.

## Acknowledgements

This work was supported by RO1 EY022030-08, RO1 EY032141-01 (AJF) and RO1 EY026158 and Kentucky Lions Eye Research Endowed Chair (MAM). We thank Isabella Palazzo for comments and discussions that shaped the final form of the manuscript.

---

**Supplemental Figure 1.**
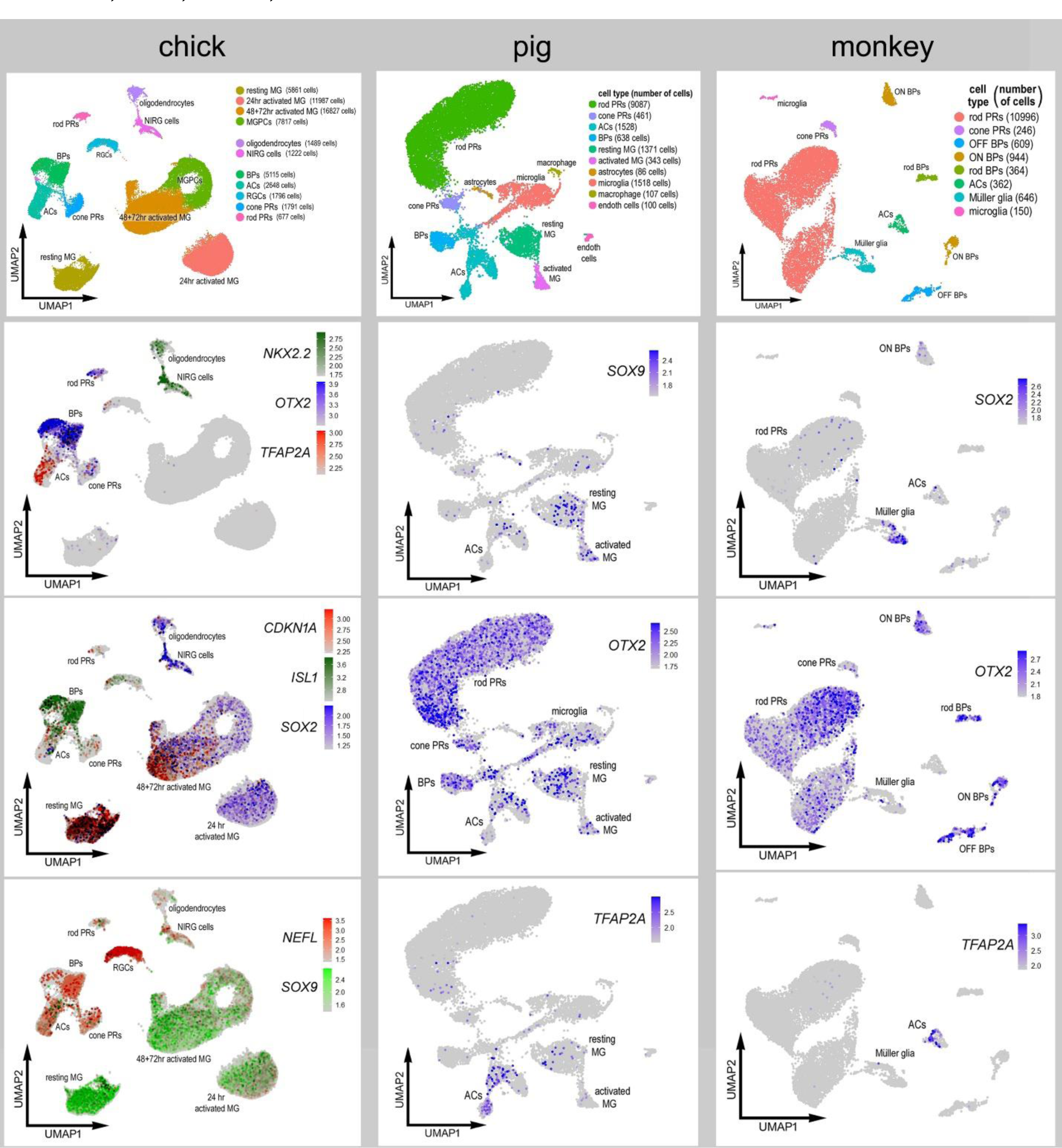
scRNA-seq libraries were probed to validate patterns of expression of markers used for immunolabeling. Retinal libraries from chick, pig and monkey were probed. UMAP heatmap plots demonstrate patterns of expression for *NKX2.2, OTX2, TFAP2A, CDKN1A, ISL1, SOX2, SOX9* and *NEFL*.

**Supplemental Figure 2.**
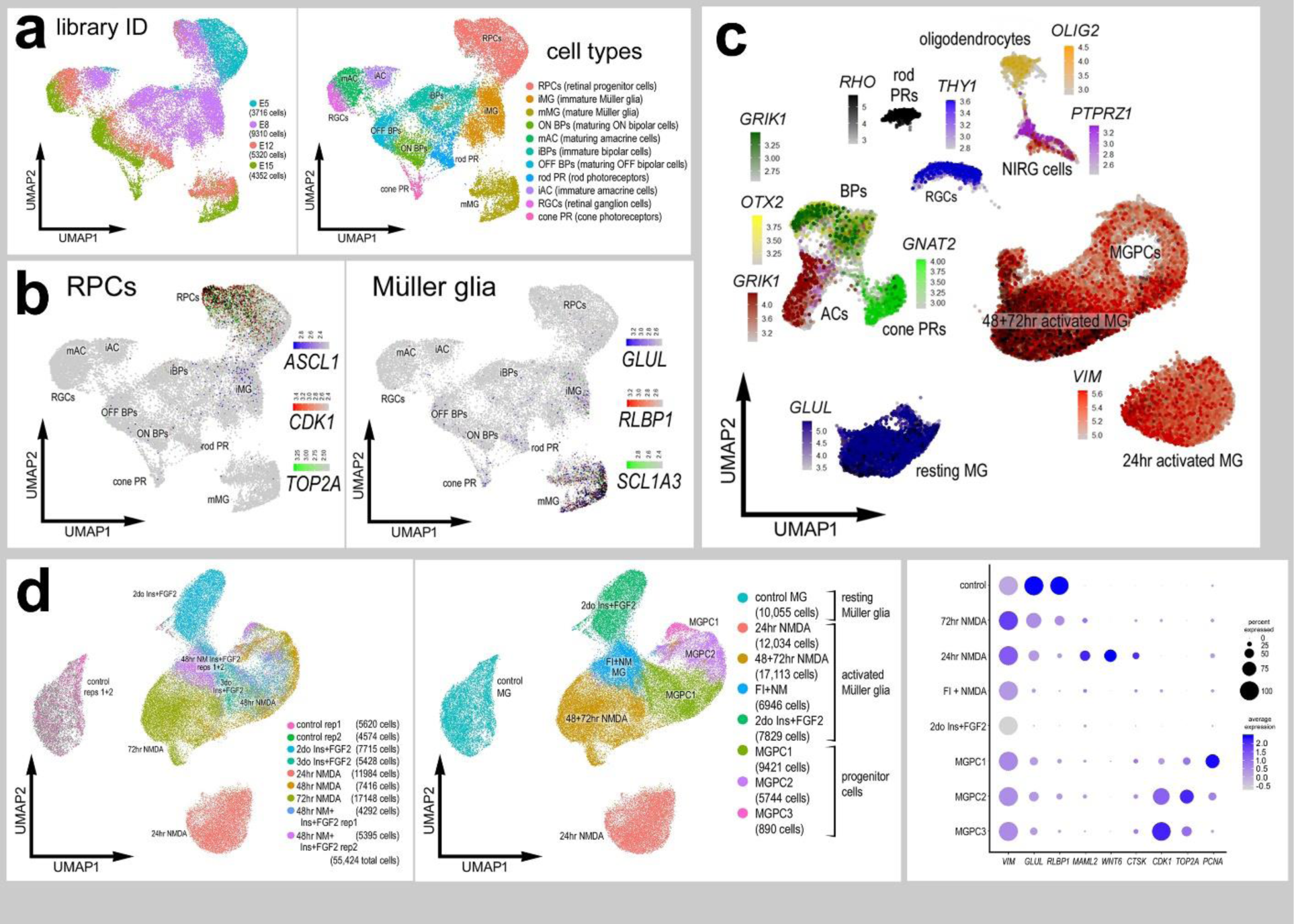
scRNA-seq libraries were generated for embryonic retinal cells at four stages of development, E5, E8, E12 and E15. UMAP-ordered clusters of cells were identified based on expression of cell-distinguishing markers, including genes characteristic of progenitors (*ASCL1, CDK1, TOP2A*) and maturing MG (*GLUL, RLBP1, SLC13A*) (a,b).(c) scRNA -libraries from post-hatch chick retinas were generated and UMAP-ordering of cells revealed distinct clusters of cells. Clusters of cells were identified using different cell-distinguishing genes including *TFAP2A* (amacrine cells), *GRIK1, OTX2* (bipolar cells), *GNAT2* (cone photoreceptors), *RHO* (rod bipolar cells), *THY1* (retinal ganglion cells), *OLIG2* (oligodendrocytes), *PTPRZ1* (NIRG cells), *GLUL* (resting MG), and *VIM* (activated MG and MGPCs). scRNA-seq was used to identify patterns of expression of NFIs in MG and MGPCs at different time points after NMDA-damage and/or treatment with FGF + insulin. Nine different libraries were aggregated for a total of more than 55,000 MG and MGPCs (d). The Dot plot illustrates the average level of expression and percentage of expressing cells resting glial genes (*VIM, GLUL, RLBP1*), genes characteristic of activated glial (*MAML2, WNT6, CTSK*), and proliferating MGPCs (*CDK1, TOP2A, PCNA*) (d).

**Supplemental Figure 3.**
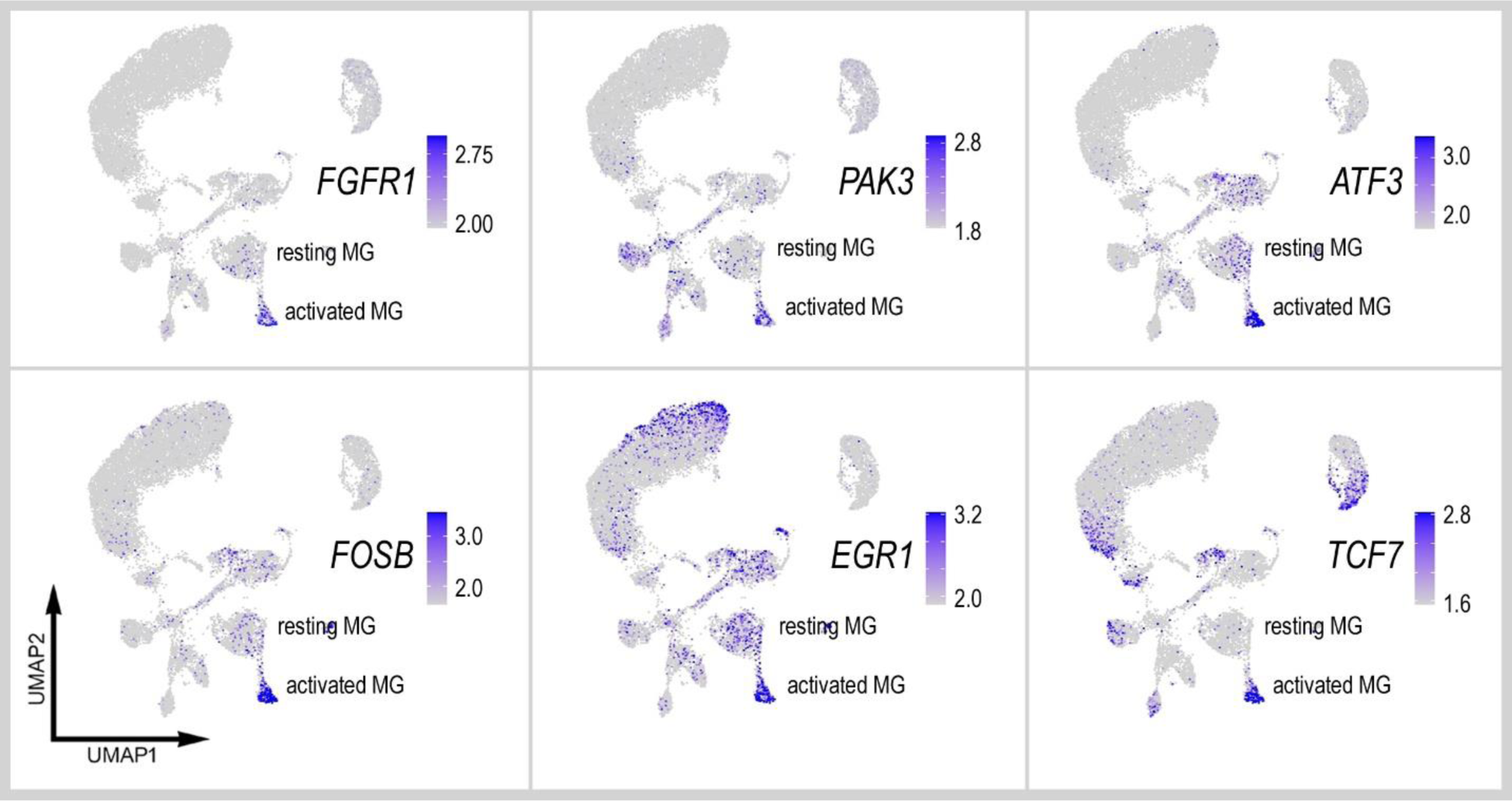
scRNA-seq was used to identify patterns of expression of genes associated with activated glia in adult pig retinas. Four different scRNA-seq libraries were aggregated for a total of 15,236 cells. UMAP clustered cells were identified based on cell-distinguishing markers (see figure 8). The expression of *FGFR1, PAK3, ATF3, FOSB, EGR1* and *TCF7* is illustrated in UMAP heatmap plots.

